# Evaluation of waste paper for cultivation of oyster mushroom (*Pleurotus ostreatus*) with some added supplementary materials

**DOI:** 10.1101/694117

**Authors:** Teklemichael Tesfay, Tesfay Godifey, Roman Mesfin, Girmay Kalayu

**Affiliations:** Department of Biology, Aksum University, Tigray, Ethiopia

**Keywords:** Biological efficiency, Oyster Mushroom, *Pleurotus ostreatus*, spawn, waste paper

## Abstract

Mushroom cultivation is an economically feasible bio-technological process for conversion of various lignocellulosic wastes. This study was conducted at Aksum University with the aim of evaluating the suitability of waste paper supplemented with corn stalk and wheat bran for Oyster mushroom cultivation. Spawn were prepared in Microbiology laboratory and inoculated into the prepared substrates. Waste paper supplemented with corn stalk and wheat bran with 0%, 25% and 50% were tested for their productivity and biological efficiency (BE) for cultivation of *P.ostreatus* mushroom. Data were analyzed using SPSS version 20. Higher (26.20± 19.36) mean weight, pileus diameter (7.90 ±2.66cm), total yield (646.4 ±273.1gm) and BE (64.64± 273 % were obtained from waste paper (50%) +cornstalk (25%) +wheat bran (25%). However, Lower (17.92±81.95%) BE were obtained from waste paper (100%). Moreover, the highest (3.88 ±0.32 cm) mean stalk length were obtained from waste paper (50%) + cornstalk (50). This study revealed that waste paper supplemented with corn stalk and wheat bran results in high BE and total yield. Thus, appears to be a promising alternative for the cultivation of oyster mushroom. Yet, waste paper without supplement poorly supports the growth of *P.ostreatus* mushroom.

## 1. Introduction

Mushrooms are fleshy, spore-bearing, multicellular fungi. They fall under the phyla Basidiomycota. Mushrooms are a good source of protein, vitamins and minerals and are known to have a broad range of uses both as food and medicine. Oyster mushroom, *Pleurotus ostreatus*, has been widely cultivated and commercialized next to *Agaricus bisporus*. Several studies have reported that *P. ostreatus* contains approximately 100 bioactive compounds, which is a potential source of dietary fiber. Besides, they are rich in protein, lipids, carbohydrates, vitamin and minerals content but low in calories and fat content (Deepalakshmi and Mirunalini, 2014). They are the easiest and least expensive commercial mushrooms to grow because they are well known for conversion of crop residues to food protein and are considered as potential source of income, alternative food production, provision of employment, and for recycling of agricultural wastes (Banik and Nandi, 2004).

Oyster mushroom has abilities to grow at a wide range of temperatures utilizing various lignocelluloses (Sánchez, 2010). Oyster mushrooms produce extensive enzymes and utilize complex organic compounds which occur as agricultural wastes and industrial by-products (Baysal *et al*., 2003). Thus, most organic matters containing cellulose, hemicellulose and lignin can be used as mushroom substrate i.e. rice and wheat straw, cottonseed hulls, corncob, paddy straw sugarcane baggase, sawdust, waste paper, and leaves (Sharma *et al*., 2013). However, an ideal substrate should contain nitrogen (supplement) and carbohydrates for rapid mushroom growth (Khare *et al.*, 2010). Oyster mushroom cultivation can play an important role in managing organic wastes, such as Waste papers and cornstalks, whose disposal has become a problem. Therefore, the current study was aimed at evaluating waste paper supplemented with cornstalk, and wheat bran as substrates for the cultivation of mushroom.

## 2. METHODOLOGY

### 2.1. DESCRIPTION OF THE STUDY AREA

The study was conducted in Microbiology laboratory, Department of Biology, Aksum University. Aksum University is found in Aksum town, 1024 km to north of Addis Ababa, Ethiopia.

### 2.2 SPAWN PREPARATION

For spawn preparation, 15 kg of sorghum was soaked in water overnight. The excess water was drained off and (5%) wheat bran and (2%) gypsum were added. The ingredients were thoroughly mixed and moisture was adjusted to 55-60%. Then the mixture was distributed equally in to 250 ml plastic bags, at the rate of 250 g seed per plastic bag and autoclaved, at 121 °C for 30 min. After cooling, each bottle was inoculated with fungal culture. When the mixture was totally invaded by mycelium, after 15 days of incubation at 25 °C, the spawn was ready to be used for the inoculation of the solid substrate (Fan *et al*., 2000).

### 2.3 Oyster Mushroom Cultivation Techniques

Oyster mushroom cultivation was done according to (Randive, 2012). The compositions of substrates used as a treatment groups for the cultivation of oyster mushroom were:

| S/NO. | Treatment groups |
| --- | --- |
| 1. | Waste paper (50%) + wheat bran (50%) |
| 2 | Waste paper 75% + wheat bran 25% |
| 3 | Waste paper 100% |
| 4 | Waste paper 50% + cornstalk 50% |
| 5 | Waste paper 75%+ corn stalk 25% |
| 6 | Waste paper 50%+corn stalk 25%+ wheat bran 25%. |

Initially, waste paper and cornstalk were chopped into small pieces (3–5 cm long). The substrates were soaked in water for 24 hours to moisten it thoroughly and pasteurized using clean steel drums. First the water was heated at 60ºC. Then the substrate was added and allowed to remain in the water for 30 minutes. Finally, once pasteurized, it was stalked on the steep cemented floor so as to remove the excessive moisture from the substrates to get 65-75% moisture level. Holes were prepared for aeration in the 500ml plastic bag. Eventually the spawn prepared was mixed with substrate and placed in dark room in 500 ml plastic bags. After the spawn run, the bags of mycelial colonized substrates were transferred to the cropping room, a room with a limited light, and watered periodically.

### 2.4. Harvesting and yield measures

Mature mushrooms were picked by clean hand without harming the substrate when they started to wrinkle-ripe. This was done for three subsequent flushes. Following the method of Iqbal *et al*. (2005), the yield parameters were recorded with respect to time (days) taken for completion of spawn running, time taken for the first appearance of pinhead formation, time taken for maturity of fruit bodies, number of flushes, and yield of flushes on the treatment substrates (Total weight of all the fruiting bodies harvested from all the three pickings were measured and considered as total yield of mushroom). The pileus diameter and the stipe length were measured with graduated transparent ruler. Mature mushrooms were weighed with analytical balance to determine the biological efficiency (BE) of mushrooms produced from substrates. The average Biological efficiency (BE) of harvests was computed as per described by Peng *et al*. (2000).

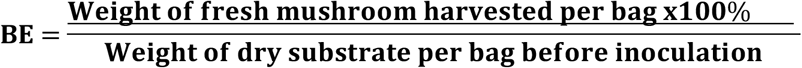

**Days Required for Completing Mycelium Running:** Time taken Days required from inoculation to completion of mycelium running was measured.

**Primordia Initiation (days):** Time required from stimulation to primordia initiation (days) were recorded.

**Number of total primordial:** Total numbers of primordial were counted from each plastic bag.

**Time from Primordial Initiation to Harvest (Maturity) (days):** Time required from primordial initiation to harvest (days) were recorded.

**Number of flushes:** The numbers of flushes were counted in each plastic bag

**Average Weight of Individual Fruiting Body/plastic bag:** Average weight of individual fruiting body was calculated by dividing the total weight of fruiting body per plastic bag by the total number of fruiting body per plastic bag. i.e.

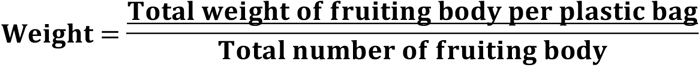

**Average Number of Effective Fruiting Body/Plastic bag:** Number of very well-developed fruiting body was recorded. Tiny fruiting bodies were discarded from counting.

**Pileus thickness (cm**): of the three randomly selected fruiting bodies of fresh mushroom pileus thickness was measured using a string.

**Mushroom pileus diameter:** The mushroom pileus diameter was taken from one end of the pileus to the other passing through the centre of the pileus and measured in millimeters (mm). This was done using a string which was placed along a ruler to get the diameter. The pileus diameter was obtained on 3 randomly picked mushrooms, from the harvest and then the average pileus diameter was calculated for a given harvest.

**Mushroom stipe length:** Stipe length was taken on the three mushrooms chosen to take the pileus diameter, using a string. The length was measured by placing the string from one end where it was attached to the substrate to the point where the gills on the pileus start on the stipe. The string was placed along a ruler to get the length in millimeters (mm).

**Yield of mushroom**= Total weight of all the fruiting bodies harvested from all the three pickings were measured as total yield of mushroom.

Five replicas of each growing trial were performed. The data on spawn running was recorded after complete colonization of substrate and pin head and fruit body formation were observed.

## 3. DATA COLLECTION AND DATA ANALYSIS

Data on mycelium colonization period, pin head formation, stalk length, BE, step length, pileus diameter were recorded and analyzed using SPSS. Analysis of variance (ANOVA) was used to indicate significant mean differences at 95% confidence interval.

## 4. RESULT

The number of days taken for complete mycelial growth differs significantly among the treatments. In the current study, the fastest mycelia extension was observed in treatment one (15 days), treatment three (15 days), and treatment five (15 days) (Table 1). Treatment 2 and Treatment 4 took the maximum numbers of days (21 and 17) respectively.

**Table 1:** The effect of substrates on mycelium colonization period (days)

| Substrate | Mean ± S.D | | | 95% | |
| --- | --- | --- | --- | --- | --- |
| | Flush1 | Flush2 | Flush3 | Confidence for<br>the Overall<br>mean | Over all<br>mean ± S.D |
| WP (50%) + CS (50%) | 15<sup>a</sup> | 15<sup>a</sup> | 15<sup>a</sup> | 15,15 | 15<sup>a</sup> |
| WP (75%) + CS (25%) | 21 ± 1.6<sup>b</sup> | 21 ± 1.6<sup>b</sup> | 21 ± 1.6<sup>b</sup> | 20.11,21.88 | 21 ± 1.6<sup>b</sup> |
| WP (50%) + CS (25%) +WB (25%) | 15<sup>a</sup> | 15<sup>a</sup> | 15<sup>a</sup> | 15,15 | 15<sup>a</sup> |
| WP 100% | 17<sup>a</sup> | 17<sup>a</sup> | 17<sup>a</sup> | 17,17 | 17<sup>ac</sup> |
| WP (75%) + WB (25%) | 15<sup>a</sup> | 15<sup>a</sup> | 15<sup>a</sup> | 15,15 | 15<sup>a</sup> |
| Cotton husk (Control) :T6 | 15<sup>a</sup> | 15<sup>a</sup> | 15<sup>a</sup> | 15,15 | 15<sup>a</sup> |

Means followed by different superscript letters within a row and columns are significantly different ( $p < 0.05$ using Tukey's multiple comparisons method), WP= waste paper, CS= Corn stalk, WB= Wheat barn

Table 2 shows that the mean pin head formation of some of the treatments varies significantly (P>0.05). Moreover, there is variation in pin head formation between flushes of each treatment. Time taken for initial appearance of pinhead after spawning of the substrate were 9.46±0.8 and 11.60 ±3.24 days for treatment group three and five respectively. Thus, treatment three and five has shown a better substrate in case of pin-head formation.

**Table 2:** The effect of substrates on pin head formation

| Substrate | mean ± SD and 95% CI of Pin for each flush | | | 95% CI for | Over all mean |
| --- | --- | --- | --- | --- | --- |
| | flush 1 | flush 2 | flush 3 | the Overall | Mean ± SD |
| WP (50%) + CS | $13 \pm 1.22^a$ | $18.2 \pm 1.30^b$ | $16.2 \pm 1.09^{bc}$ | 14.42, 17.74 | $15.8 \pm 2.48^a$ |
| (50%) | 11.47, 14.12 | 16.58, 19.82 | 14.84, 17.56 |  |  |
| WP (75%) + CS | $12.8 \pm 0.84^a$ | $15.60 \pm 2.30^a$ | $17.20 \pm 2.16^a$ | 13.77, 16.62 | $15.20 \pm 2.56^a$ |
| (25%) | 11.76, 13.84 | 12.74, 18.46 | 14.50, 19.89 |  |  |
| WP (50%) + CS | $5.80 \pm 1.30^a$ | $10.80 \pm 0.86^b$ | $11.80 \pm 1.64^{bc}$ | 7.74, 11.19 | $9.46 \pm 0.80^b$ |
| (25%) + WB (25%) | 4.18, 7.42 | 8.41, 13.18 | 9.76, 13.84 |  |  |
| WP 100% | $19 \pm 2.34^a$ | $16.20 \pm 5.67^a$ | $14.40 \pm 3.20^a$ | 14.21, 18.85 | $16.53 \pm 4.18^a$ |
|  | 16.08, 21.91 | 9.15, 23.24 | 10.41, 18.38 |  |  |
| WP (75%) + WB | $14.60 \pm 2.79^a$ | $9.80 \pm 3.03^b$ | $10.40 \pm 1.67^{ac}$ | 9.80, 13.39 | $11.60 \pm 3.24^c$ |
| (25%) | 11.13, 18.06 | 6.03, 13.56 | 8.32, 12.47 |  |  |
| Control | 8 | 10 | 9 | 6.51, 11.48 | $9.0 \pm 1.00^d$ |

Means followed by different superscript letters within a row and columns are significantly different ( $p < 0.05$ using Tukey's multiple comparisons method), WP= waste paper, CS= Corn stalk, WB= Wheat barn

Table 3 indicates mean ±SD for each flush and the overall maturity (days) of Oyster mushroom. Maturity were not significantly different (p>0.05) among the flush of each treatment while among the treatments, treatment four (Waste paper 100%) were significantly (p<0.05) different. Considering the minimum number of days taken for maturity of fruiting bodies, treatment one (3.4, 3.6 and 3.4 days) appears to be the best substrate followed by treatment three (4.2,3.6 and 3.2 days) (Table 3). Maximum time period (4.4, 4.4,4 days) was required for the maturity of fruiting bodies in case of treatment four (waste paper (100%). Besides maturity between treatments were not significantly (p>0.05) different. The mean maturity of the different treatments ranges from 3.47 ±0.52 (Treatment 1) to 4.27± 0.88 (Treatment 4). However, it took less days for maturation compared to the control group.

**Table 3:** Period of pinning-to-maturation of mushrooms in substrates (Days)

| Substrate | mean ± SD and 95% CI of Maturity for each flush | | | 95% CI for the | Over all mean |
| --- | --- | --- | --- | --- | --- |
| | flush 1 | flush 2 | flush 3 | Overall mean | Mean ± SD |
| WP (50%) + CS (50%) | 3.40 ± 0.55<sup>a</sup> | 3.60 ± 0.55<sup>a</sup> | 3.40 ± 0.548<sup>a</sup> | 3.18 ,3.75 | 3.47 ± 0.52<sup>a</sup> |
|  | 2.72 , 4.08 | 2.92 , 4.28 | 2.72 , 4.08 |  |  |
| WP (75%) + CS (25%) | 3.60 ± 0.55<sup>a</sup> | 3.80 ± 0.45<sup>a</sup> | 3.80 ± 0.45<sup>a</sup> | 3.48 ,3.99 | 3.73 ± 0.46<sup>a</sup> |
|  | 2.92 ,4.28 | 3.24 , 4.36 | 3.24 , 4.36 |  |  |
| WP (50%) + CS (25%)<br>+WB (25%) | 4.20 ± 0.84<sup>a</sup> | 3.60 ± 0.55<sup>a</sup> | 3.20 ± 0.45<sup>a</sup> | 3.27 , 4.07 | 3.67 ± 0.72<sup>a</sup> |
|  | 3.16 , 5.24 | 2.92 , 4.28 | 2.64 , 3.76 |  |  |
| WP 100% | 4.40 ± 0.89<sup>a</sup> | 4.40 ± 0.89<sup>a</sup> | 4.00 ± 1.00<sup>a</sup> | 3.78 , 4.76 | 4.27 ± 0.88<sup>b</sup> |
| | 3.29 , 5.51 | 3.29 , 5.51 | 2.76 ± 5.24 | | |
| WP (75%) + WB (25%) | 4.00 ± 0.71<sup>a</sup> | 3.60 ± 0.55<sup>a</sup> | 3.60 ± 0.55<sup>a</sup> | 3.40 , 4.76 | 3.73 ± 0.59<sup>a</sup> |
|  | 3.12 , 4.88 | 2.92 , 4.28 | 2.92 , 4.28 |  |  |
| Control | 4<sup>a</sup> | 4<sup>a</sup> | 6<sup>a</sup> | 1.34 , 4.06 | 4.53 ± 0.74<sup>a</sup> |

Means followed by different superscript letters within a row and columns are significantly different (p<0.05 using Tukes multiple comparisons method), WP= waste paper, CS=Corne stalk, WB= Wheat barn

Table 4 indicates that higher mean stalk length were measured in treatment one (3.88 0.32 cm) followed by treatment three (3.62 0.36 cm). However, no significant difference was observed in terms of stalk length between the different substrates and the control. But, a decreasing pattern was observed in terms of stalk length of flush in each treatment.

**Table 4:**
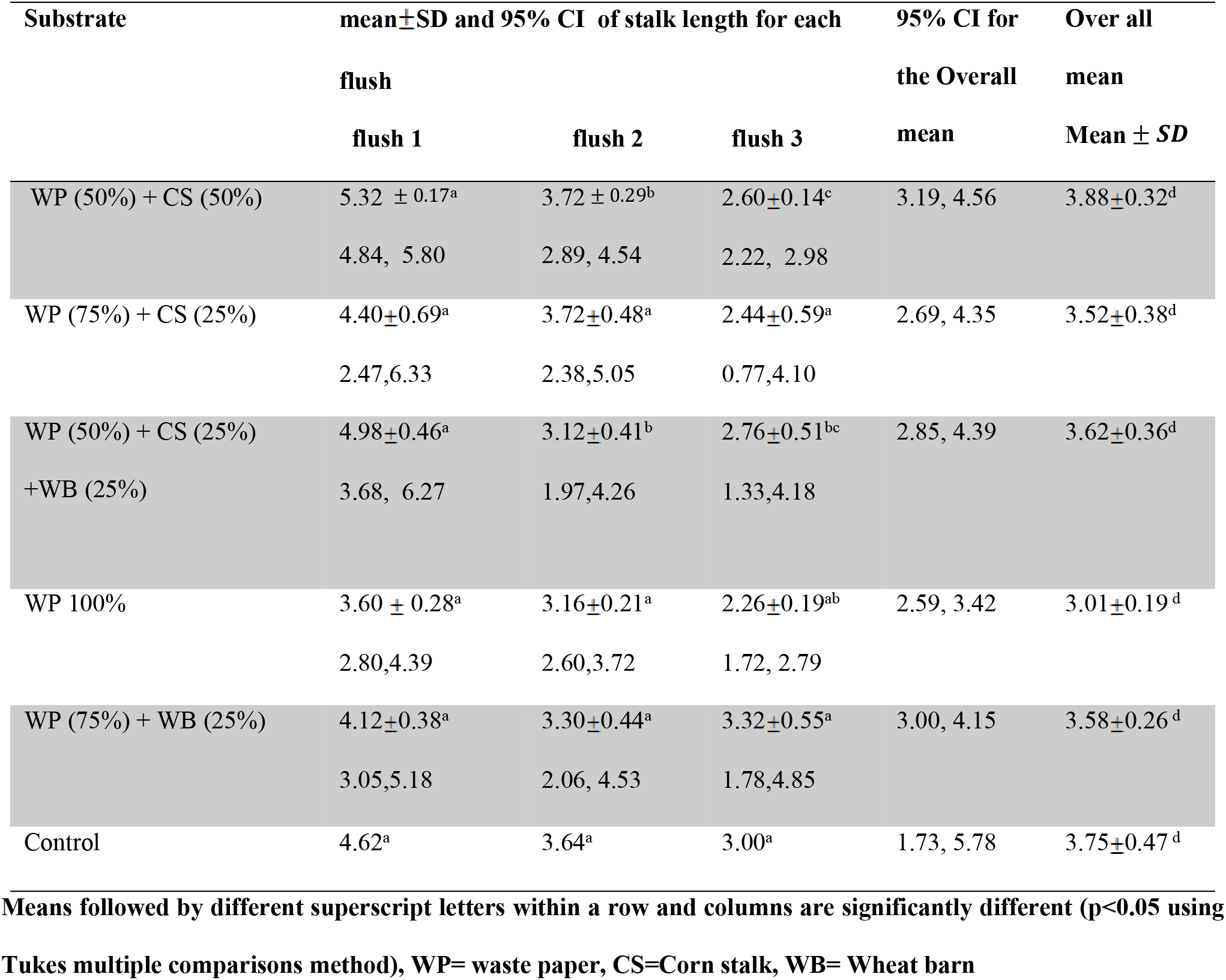
The effect of substrate on stalk length (cm)

| Substrate | mean ± SD and 95% CI of stalk length for each flush | | | 95% CI for the | Over all |
| --- | --- | --- | --- | --- | --- |
|  | flush 1 | flush 2 | flush 3 | the Overall mean | mean |
| WP (50%) + CS (50%) | 5.32 ± 0.17<sup>a</sup> | 3.72 ± 0.29<sup>b</sup> | 2.60±0.14<sup>c</sup> | 3.19, 4.56 | 3.88±0.32<sup>d</sup> |
|  | 4.84, 5.80 | 2.89, 4.54 | 2.22, 2.98 |  |  |
| WP (75%) + CS (25%) | 4.40±0.69<sup>a</sup> | 3.72±0.48<sup>a</sup> | 2.44±0.59<sup>a</sup> | 2.69, 4.35 | 3.52±0.38<sup>d</sup> |
|  | 2.47,6.33 | 2.38,5.05 | 0.77,4.10 |  |  |
| WP (50%) + CS (25%)<br>+WB (25%) | 4.98±0.46<sup>a</sup> | 3.12±0.41<sup>b</sup> | 2.76±0.51<sup>bc</sup> | 2.85, 4.39 | 3.62±0.36<sup>d</sup> |
|  | 3.68, 6.27 | 1.97,4.26 | 1.33,4.18 |  |  |
| WP 100% | 3.60 ± 0.28<sup>a</sup> | 3.16±0.21<sup>a</sup> | 2.26±0.19<sup>ab</sup> | 2.59, 3.42 | 3.01±0.19<sup>d</sup> |
|  | 2.80,4.39 | 2.60,3.72 | 1.72, 2.79 |  |  |
| WP (75%) + WB (25%) | 4.12±0.38<sup>a</sup> | 3.30±0.44<sup>a</sup> | 3.32±0.55<sup>a</sup> | 3.00, 4.15 | 3.58±0.26<sup>d</sup> |
|  | 3.05,5.18 | 2.06, 4.53 | 1.78,4.85 |  |  |
| Control | 4.62<sup>a</sup> | 3.64<sup>a</sup> | 3.00<sup>a</sup> | 1.73, 5.78 | 3.75±0.47<sup>d</sup> |

Means followed by different superscript letters within a row and columns are significantly different ( $p < 0.05$ using Tukey's multiple comparisons method), WP= waste paper, CS=Corn stalk, WB= Wheat barn

Table 5 shows that the mean ±SD of each flush and the overall mean of pileus diameter. The highest (7.90 ± 2.66 cm) and the lowest (5.40 ± 1.57cm) mean pileus diameter were noted on treatment three and five respectively. Significant difference was observed between treatments 3 and 4. Besides, pileus diameter among flushes were not significantly (p<0.05) different except in the second and third flushes of treatments two and three (Table 5).

**Table 5:**
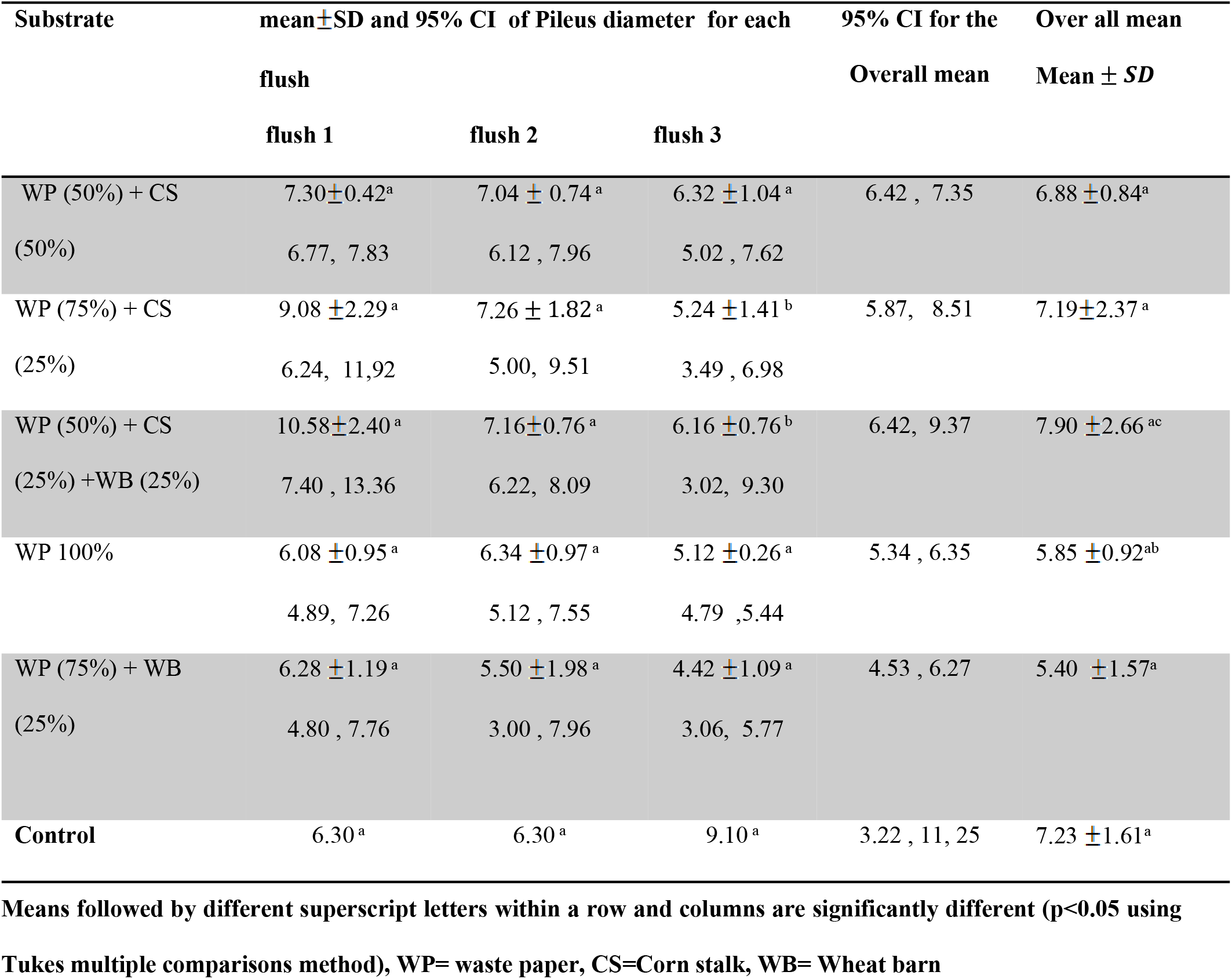
The effect of substrates on Pileus diameter (cm)

| Substrate | mean ± SD and 95% CI of Pileus diameter for each flush | | | 95% CI for the Overall mean | Over all mean Mean ± SD |
| --- | --- | --- | --- | --- | --- |
|  | flush 1 | flush 2 | flush 3 |  |  |
| WP (50%) + CS | 7.30±0.42<sup>a</sup> | 7.04 ± 0.74<sup>a</sup> | 6.32 ± 1.04<sup>a</sup> | 6.42 , 7.35 | 6.88 ± 0.84<sup>a</sup> |
| (50%) | 6.77, 7.83 | 6.12 , 7.96 | 5.02 , 7.62 |  |  |
| WP (75%) + CS | 9.08 ± 2.29<sup>a</sup> | 7.26 ± 1.82<sup>a</sup> | 5.24 ± 1.41<sup>b</sup> | 5.87, 8.51 | 7.19±2.37<sup>a</sup> |
| (25%) | 6.24, 11.92 | 5.00, 9.51 | 3.49 , 6.98 |  |  |
| WP (50%) + CS | 10.58±2.40<sup>a</sup> | 7.16±0.76<sup>a</sup> | 6.16 ± 0.76<sup>b</sup> | 6.42, 9.37 | 7.90 ± 2.66<sup>ac</sup> |
| (25%) +WB (25%) | 7.40 , 13.36 | 6.22, 8.09 | 3.02, 9.30 |  |  |
| WP 100% | 6.08 ± 0.95<sup>a</sup> | 6.34 ± 0.97<sup>a</sup> | 5.12 ± 0.26<sup>a</sup> | 5.34 , 6.35 | 5.85 ± 0.92<sup>ab</sup> |
|  | 4.89, 7.26 | 5.12 , 7.55 | 4.79 , 5.44 |  |  |
| WP (75%) + WB | 6.28 ± 1.19<sup>a</sup> | 5.50 ± 1.98<sup>a</sup> | 4.42 ± 1.09<sup>a</sup> | 4.53 , 6.27 | 5.40 ± 1.57<sup>a</sup> |
| (25%) | 4.80 , 7.76 | 3.00 , 7.96 | 3.06, 5.77 |  |  |
| Control | 6.30<sup>a</sup> | 6.30<sup>a</sup> | 9.10<sup>a</sup> | 3.22 , 11, 25 | 7.23 ± 1.61<sup>a</sup> |

Means followed by different superscript letters within a row and columns are significantly different ( $p < 0.05$ using Tukes multiple comparisons method), WP= waste paper, CS= Corn stalk, WB= Wheat barn

Table 6 indicates the effect of substrate on mushroom Weight (gm). Of the 1^st^ flush generation, the maximum (34.08 ± 45.69gm) and minimum (6.34 ± 1.44 gm) mean weight gm) were recorded on treatment two and one respectively. Of the 2^nd^ flush generation, the highest mean weight (33.76 ± 22.47) was recorded on treatment four. Mean weight of harvested flush decrease with successive generations (Table 6). Besides, the higher (26.20 ± 19) value of overall mean weight of individual fruiting body was observed in treatment three.

**Table 6:** The effect of substrates on Weight (gm)

| Substrate | mean ± SD and 95% CI of Weight for each flush | | | 95% CI for the | Over all mean |
| --- | --- | --- | --- | --- | --- |
| | flush 1 | flush 2 | flush 3 | Overall mean | Mean ± SD |
| WP (50%) + CS (50%) | $6.34 \pm 1.44^a$<br>4.54, 8.14 | $4.94 \pm 1.71^a$<br>2.82, 7.06 | $5.08 \pm 2.31^a$<br>2.21, 7.95 | 4.43, 6.47 | $5.45 \pm 1.84^a$ |
| WP (75%) + CS (25%) | $34.08 \pm 45.69^a$<br>-22.66, 90.82 | $14.04 \pm 12.38^a$<br>-1.34, 29.42 | $15.36 \pm 4.97^a$<br>9.18, 21.54 | 6.12, 36.19 | $21 \pm 27.15^a$ |
| WP (50%) + CS (25%)<br>+WB (25%) | $33.90 \pm 1.06^a$<br>21.40, 46.39 | $32 \pm 24.92^a$<br>1.05, 62.94 | $12.70 \pm 15.71^a$<br>-6.80, 32.21 | 15.47, 36.93 | $26.20 \pm 19.36^b$ |
| WP 100% | $21.16 \pm 11.15^a$<br>7.31, 35.05 | $33.76 \pm 22.47^a$<br>5.85, 61.66 | $14.56 \pm 11.00^a$<br>0.89, 28.22 | 13.85, 32.48 | $23.16 \pm 16.80^a$ |
| WP (75%) + WB (25%) | $28.20 \pm 7.46^a$<br>18.93, 37.47 | $24.64 \pm 20.16^a$<br>-0.39, 49.67 | $21.38 \pm 13.38^a$<br>4.76, 37.99 | 17.07, 32.40 | $24.74 \pm 13.84^b$ |
| Control | $27.4^a$ | $19.7^a$ | $21.33^a$ | 12.71, 32.89 | $22.80 \pm 4.06^a$ |

Means followed by different superscript letters within a row and columns are significantly different ( $p < 0.05$ using Tukes multiple comparisons method), WP= waste paper, CS= Corn stalk, WB= Wheat barn

Table 7 indicates the effect of the treatment groups with varying substrate composition on yield (gm) and BE (%). The highest total yield (682.1gm) was obtained from control followed by treatment three (646.4 ± 273.1 gm). Of the 1^st^ flush cropped, the maximum yield (435.86 ± 133.34 gm) was recorded on treatment three while the lowest (87.4 ±48.07) yield was obtained from treatment four. On the other hand, in 2^nd^ generation flush the mean yield ranged from 57.40± 15.85 (gm) to 232 gm and highest was recorded on cotton husk (control). In the 3^rd^ generation flush the minimum (34.40 ± 18.06g) total yield was recorded on treatment four. Though, significant (p>0.05) difference was observed only in treatment four ignoring the flushes.

**Table 7:**
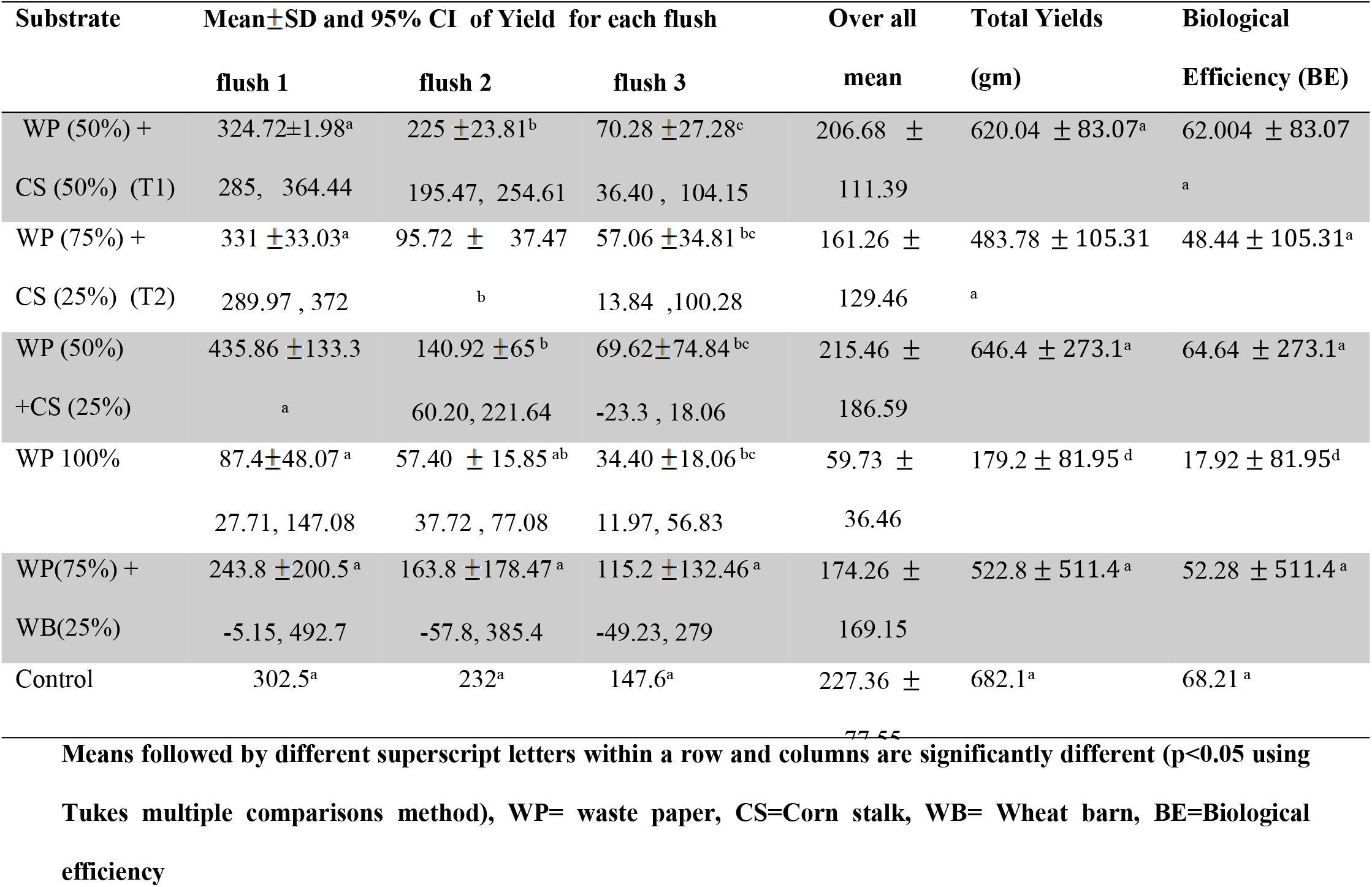
The effect of substrate on yield (gm) and BE (%)

## 5. DISCUSSION

Mycelial growth provides suitable internal conditions for fruiting. In this study, the fastest mycelia extension was observed in treatment one (15 days), three, and five equally. Thus, outstanding growth of mycelium is a vital factor in mushroom cultivation (Pokhrel *et al*., 2009). In this study, waste paper supplemented with wheat bran and corn stalk produce mycelium extension within short period of time which is similar with the control except the second treatment group. However, waste paper without supplementary materials takes relatively rather extended time. This could be due to the variation in nutrient content, lignin and cellulose composition and moisture holding capacity of the substrate. Similar results were reported by Shah *et al.*, (2004) where the growth of *Pleurotus* species on wheat straw, rice husk as well as saw dust took 2-3 weeks for spawn running (mycelial growth) after inoculation. Moreover, Kumari and Achal (2008) noted that colonization of the substrate with *P.ostreatus* was completed within 20 days of inoculation. Conversely, the current study contradicts with the results of Zenebe Girmay *et.al* (2016) where they reported that mycelia running in waste paper took 14 days. The variation in mycelia extension might be due to the difference in condition of the environment and the nature of the substrate. *P. ostreatus* grew quickly at 30 °C (Marino *et al*., 2003) and oyster yield decreases when the temperature decreases in different climatic zones (Zervakis *et al*., 2001).

Sharma *et al* (2013) reported that primordial initiation (pin head formation) on various substrates were in between 26.40-31.60 days of incubation. Moreover, Shah *et al*. (2004) indicated that relatively higher room temperature could have resulted in shorter pining periods (27 to 34 days of incubation). This contradicts with our result where all the treatments initiate the pin head within few days. Oei (2003) reported that materials with high quality lignin and cellulose contents take longer time to start pinning compared to the substrates with low contents of the lignin and cellulose. This study reveals that as the amount of waste paper increases the time taken for pinning increases (Table 2). Thus, the longer time taken for pinning might be due to the cellulose and lignin content of waste paper. Different scholars reported different pinning days. The variation in pin head formation might be due to the difference in room temperature of the cultivation room and nutrient availability of the substrate.

A number of investigators have reported different timing period for fruiting bodies (maturity). Similar results (4 ± 0.7 days) for the maturation of fruit bodies were reported by Gume *et al.*, (2013). The current result is inconsistent with Islam *et al*. (2009) which reported that maturation period of *Pleurotus* species ranging from 3.29 to 4.33 growing on saw dusts of Mango, Shiris, Jackfruit, Kadom, Jam and Coconut. Moreover, Girmay *et.al* (2016) noted a higher (39) number of maturation days of *P.ostreatus* mushroom cultivated on waste paper. The variation in maturity of fruiting bodies could be owing to the difference in physiological requirements and the nature of the substrate.

Gume *et al*. (2013) reported shorter (1.4 to1.9 cm) stalk length and pileus diameter (3.8 to 5.2 cm) than the current finding on mushrooms grown on sawdust, coffee bean husks, and corncobs. Stipe (stalk) length and pileus diameter of oyster mushroom grown in different substrates depended on the structure, compactness and physical properties of the substrate which in turn depend on the type of substrates. The substrates with higher moisture retaining capacity perform better than those with lower moisture retaining capacity (Chukwurah *et. al*., 2013). Fruit bodies with larger pileus (caps) and shorter stipes (stalk) are better than that with smaller pileus and longer stipes (Synytsya *et al.*, 2008). In the current study, treatment two provides better quality of mushroom with larger pileus diameter and shorter stalk length. However, the stipes contains more insoluble dietary fibers that can be used for the preparation of biologically active polysaccharide complexes utilizable as food supplements than pilei. Moreover, Kivaisi *et.al* (2003) indicated that the size of the pileus depends on the aeration and amount of light.

Sarker *et al.* (2007) reported that the individual weight of fruiting body ranged from 1.33-1.59 g, which was less than the current finding. Similarly Bhuyan (2008) reported less (5.02-7.01 g) result than the current study. Moreover, Bhuyan (2008) reported a significant effect of supplementation on weight of individual fruiting bodies. The variation in weight of individual fruiting bodies might be due to environmental conditions or growing season and variation in nutrient composition of the substrates.

Yield among the flushes of each treatment varies significantly (P<0.05) for some of the treatments (Table 7). Besides, the yield of all treatments did not significantly vary. This indicates that waste paper supplemented with corn stalk and wheat bran could replace cotton husk for cultivation of mushroom. In this research, higher yield were obtained compared to Sharma *et al* (2013) with 381.85 gm yield of *Pleurotus ostreatus* growing on rice straw, rice straw + wheat straw, rice straw + paper, sugarcane bagasse and sawdust of alder. Furthermore, the maximum biological efficiency (64.64 _±273.1_) was recorded on treatment three while the lowest (17.92 ±81.95%) BE was obtained from treatment four. This is in line with the works of Holkar and Chandra (2016) where they reported that the biological efficiencies of *P.ostreatus* growing on wheat straw range from 63.4 to 74. As per Gume *et al*., (2013), substrates that gave over 40% BE could be recommended for oyster mushrooms cultivation. Thus, the current study reveals that all the treatments except waste paper (100%) without supplement gave higher BE (Table 7). This could be due to the better availability of nitrogen, carbon and minerals from the supplements (Shah *et al.*, 2004).

## 6. CONCLUSION

This study clearly indicates that Waste paper supplemented with corn stalk and wheat bran offers higher total yield and Biological efficiency. It represents promising substrates which can serves as a basal medium for the cultivation of Oyster mushroom. This biological process revealed that the conversion of waste papers into a biomass of edible mushroom and pest that can be utilized as a fertilizer. It appears that the lignin, cellulose and hemicellulose, the active components of paper, provides a carbon sources. Thus, it is ecofriendly approach in terms of solid waste management and is also economically sound in light of food security.

## REFERENCES

1. Banik S & Nandi R. Effect of supplementation of rice straw with biogas residual slurry manure on the yield, protein and mineral contents of oyster mushroom. Industrial Crops and Products. 2004; 20:311–319.

2. Baysal E., Peker H., Yalinkilic MK., and Temiz A. Cultivation of Oyster Mushroom on waste paper with some added supplementary material. Bioresource Technology. 2003; 89:95–97.

3. Bhuyan, M.H.M.B.U. Study on Preparation of Low Cost Spawn Packets for the Production of Oyster Mushroom (*Pleurotus Ostreatus*) and its Proximate Analysis, M.S. Thesis, Department of Biochemistry, SAU, Dhaka. 2008

4. Chukwurah N. F., Eze S. C., Chiejina N. V.3., Onyeonagu C. C., Okezie C. E. A. Ugwuoke K. I., Ugwu F. S. O., Aruah C. B.1, Akobueze E. U., and Nkwonta C. G. Correlation of stipe length, pileus width and stipe girth of oyster mushroom (*Pleurotus ostreatus*) grown in different farm substrates. 2013

5. Deepalakshmi K andMirunalini S. *Pleurotus ostreatus*: an oyster mushroom with nutritional and medicinal properties. J Biochem Tech.2014; 5:718–726.

6. Fan L., Pandey A., Mohan R and Soccol C. R. Use of various coffee industry residues for cultivation of *Pleurotus ostreatus* in solid state fermentation. Acta Biotechnol.2000; 20: 41–52.

7. Gume B., Muleta D., and Abate D. Evaluation of locally available substrates for cultivation of oyster mushroom (*Pleurotus ostreatus*) in Jimma, Ethiopia. African journal of Microbiology Research.2013; 7: 2228–2237.

8. Holkar K.S and Chandra R.Comparative evaluation of five *pleurotus species* for their growth behavior and yield performance using wheat straw as a substrate. Journal of environmental Biology. 2016; 37:7–12.

9. Iqbal S.M., Rauf C.A and Sheikh M. I. Yield performance of oyster mushroom on different substrate, International Journal of Agriculture and Biology.2005; 7: 900–903.

10. Islam M.Z, Rahman M.H and Hafiz F. Cultivation of oyster mushroom (*Pleurotus flabellatus*) on different substrates. Int. J. Sustain. Crop Prod.2009; 4:45–48.

11. Khare K.B., Mutuku J.M., Achwania O.S and Otaye D.O. Production of two oyster mushrooms, *Pleurotus sajor-caju* and *P. florida* on supplemented and un-supplemented substrates, International Journal of Agriculture and Applied Sciences. 2010; 6: 4–11.

12. Kivaisi K., F.S.S. Magingo and B. Mamiro. Performance of P.flabellatus on water hyacinth shoots at two different temperature and relative humidity regimes. Tanzania J.Sc. 2003; 29:11–18.

13. Kumari D and Achal V. Effect of different substrates on the production and nonenzymatic antioxidant activity of *Pleurotus ostreatus*. Life Sci J. 2008; 5:73–76.

14. Marino RH, da Eira AF, Kuramae EE and Queiroz EC. Morphomolecular characterization of *Pleurotus ostreatus* (Jacq. Fr.) kummer strains in relation to luminosity and temperature of frutification. Scientia Agricola, 2003; 60:531–535.

15. Oei P. Mushroom cultivation-Appropriate Technology for Mushroom Growers. 3^rd^ Edn., Backhuys Publishers, Leiden, Netherlands.2003

16. Peng J.T., Lee C.M and Tsai, Y.F. Effect of rice Bran on the production of different king oyster mushroom strains during bottle cultivation. J. Agric. Res. 2000; 49:60–67.

17. Randive S.D. Cultivation and study of growth of oyster mushroom on different agricultural waste substrate and its nutrient analysis. Advances in Applied Science Research.2012; 3:1938–1949.

18. Sarker N.C., Hossain M.M., Sultana N., Mian H., Karim A.J.M.S and Amin SMR. Performance of Different Substrates on the Growth and Yield of Pleurotus ostreatus (Jacquin ex Fr.) Kummer, Bangladesh J. Mushroom.2007; 1:9–20.

19. Sánchez, C. Cultivation of *Pleurotus ostreatus* and other edible mushrooms. Appl. Microbiol. Biotechnol. 2010; 85:1321–1337.

20. Sharma S., Yadav P.R and Pokhrel P.C. Growth and Yield of Oyster mushroom (*Pleurotus ostreatus*) on different substrates. Journal on New Biological Reports. 2013; 2:3–8.

21. Shah Z.A., Ashraf M and Ishtiaq M.C. Comparative study on cultivation and yield performance of Oyster mushroom (*Pleurotus ostreatus*) on different substrates (wheat straw, Leaves and Sawdust). Pakistan J. Nutrition. 2004; 3:158–160.

22. Synytsya A, Mickova K, Jablonsky I, Slukova M and Copikova J. Mushrooms of genus *Pleurotus* as a source of dietary fibers and glucans for food supplements. Czech. J. Food Sci.2008; 26: 441–446.

23. Pokhrel CP, Yadav RKP and Ohga S. Effects of physical factors and synthetic media on mycelial growth of *Lyophyllum decastes*. Jour Ecobiotech.2009; 1:46–50.

24. Zenebe Girmay, Weldesemayat Gorems, Getachew Birhanu and Solomon Zewdie. Growth and yield performance of *Pleurotus ostreatus* (oyster mushroom) on different substrates. AMB Express.2016; 6: 87.

25. Zervakis G, Philippoussis A, Ioannidou S and Diamantopoulou P. Mycelium growth kinetics and optimal temperature conditions for the cultivation of edible mushroom species on lignocellulosic substrates. Folia microbiologica.2001; 46: 231–4.

